# TDP-43 dysfunction leads to the accumulation of cryptic transposable element-derived exons, crypTEs, in iPSC derived neurons and ALS/FTD patient tissues

**DOI:** 10.64898/2026.01.09.698641

**Authors:** Isobel Bolger, Regina Shaw, Oliver H Tam, Cláudio Gouveia Roque, Christopher A Jackson, Kathryn O’Neill, NYGC ALS Consortium, Colin Smith, Hemali Phatnani, Karthick Natarajan, Molly Gale Hammell

## Abstract

TDP-43 is an RNA and DNA binding protein that plays major roles in regulating RNA processing. In particular, TDP-43 dysfunction leads to the accumulation of cryptic splice isoforms that result from improperly spliced mRNAs. In addition to its role in regulating splicing, TDP-43 is also known to regulate the expression of transposable elements (TEs). TEs are mobile genetic elements which comprise a significant proportion of the human genome, but are normally silenced in healthy somatic cells. TEs are interspersed throughout the genome, both in gene-depleted regions and within gene introns and gene regulatory sequences. We used optimized long-read RNA sequencing assays to generate catalogs of mis-spliced and mis-expressed genes and TEs in human neurons depleted for TDP-43. In addition to known TDP-43 driven cryptic isoforms, we identified hundreds of TDP-43 dependent spliced RNAs that form cryptic gene-TE fusion events as a result of mis-splicing of TE sequences into gene transcripts. Among these TDP-43 dependent cryptic gene-TE transcripts (crypTEs), we found: TEs that provide alternate gene promoters/5’UTRs, TEs that act as cassette exons inside host gene mRNAs, as well as TEs that provide alternate transcript 3’ ends. These cryptic gene-TE fusions are predicted to induce aberrant expression of ALS relevant genes, nonsense mediated decay (NMD) products, as well as novel peptides from gene-TE fusions within the gene coding sequence. Using coupled long-read RNA (Iso-seq) and single-nucleus (snRNA-seq) profiles from postmortem ALS tissues, we further verified that many of these crypTE transcripts are enriched in frontal cortex samples from ALS donors with cognitive involvement (ALSci) and associated with altered expression of those genes in deep layer cortical excitatory neurons. In short, TDP-43 dependent crypTEs greatly expand the catalogs of TDP-43 dependent cryptic splice isoforms and represent a novel mechanism by which TE dysregulation impacts ALS.

## Introduction

Amyotrophic Lateral Sclerosis (ALS) and Frontotemporal Dementia with TDP-43 pathology (FTD-TDP) are two related neurodegenerative diseases^1–3^ whose pathological hallmark involves the accumulation of large aggregates of TDP-43 protein^4,5^. TDP-43 is an RNA and DNA binding protein that plays essential roles in regulating gene expression, mRNA splicing, and mRNA trafficking^6^. TDP-43 aggregates^7,8^ are present in the tissues of 95% of ALS patients, 50% of patients with (FTD), and 25% of Alzheimer’s Disease (AD) patients. Recent reports from several groups have identified TDP-43 pathology specific RNA splicing alterations, many of which are sufficient to induce neurotoxicity downstream of TDP-43 functional loss^9–13^. Moreover, the splicing changes that result from TDP-43 dysfunction in motor cortex tissues are largely non-overlapping with the splicing changes seen in spinal cord^14,15^, suggesting highly cell-type and tissue-specific differences in TDP-43 splicing patterns^16–18^, even between upper and lower motor neurons. While a few of these TDP-43 mediated cryptic splice isoforms have been characterized for their phenotypic impact, such as STMN2^9,13^ and UNC13A/B^11,12^, the vast majority of these cryptic transcripts remain largely uncharacterized^19^.

In addition to the splicing targets of TDP-43 protein, TDP-43 is also known to regulate the expression of transposable elements (TEs)^20^. TEs are viral-like mobile genetic elements that normally lie dormant in our genomes but can become reactivated in the context of aging^21–24^ and neurodegenerative disease^14,20,25–30^. Loss of normal nuclear TDP-43 protein results in the aberrant upregulation of TEs due to the failure of TDP-43 to bind and silence TE expressed RNAs. Analysis of post-mortem cortical and spinal cord tissues from ALS/FTD patients demonstrated the clinical relevance of this role for TDP-43 in regulating TE expression, demonstrating tight correlation between elevated TE expression, TDP-43 mis-splicing events, and dense TDP-43 pathology in ALS/FTD patient tissues^14,20^.

We generated optimized protocols for long-read RNA sequencing in order to profile TDP-43 dependent changes in splice isoform usage and TE expression in neuronal-like cells depleted for TDP-43 protein. These enhanced long read RNA sequencing protocols returned hundreds of novel cryptic splice isoforms and expressed intact TEs. In addition to full-length TE and mRNA transcripts, these long-read RNA sequencing profiles also returned thousands of cryptic splice isoforms that involved fusions of TE sequence with gene mRNAs. This process of TE exonization^31,32^ has been shown to occur at low levels in normal and cancer cells. Using CRISPRi mediated depletion of TDP-43 in induced neuronal-like cells (i3Neurons), we show that TDP-43 normally prevents TE exonization to ensure splicing fidelity of its target genes. Here, we present a catalog of TDP-43 dependent cryptic TE-derived splice isoforms (crypTEs), alongside optimized protocols for cryptic gene and TE isoform sequencing (CrypTE-seq). We further characterize a set of crypTE splice isoforms that show TDP-43 dependence in cultured iPSC-derived excitatory neurons, presence in ALS/FTD cortical tissues, associations with clinical features such as cognitive impairment, and associations with alteration in gene splicing and expression in ALS/FTD cortical excitatory neurons.

## Results

### CRISPRi mediated knockdown of TDP-43 in iPSC derived neurons (i3Neurons) results in stable depletion of TDP-43 mRNA and protein

We first set out to establish a stable model of TDP-43 loss of function in differentiated neuronal-like cells that mimic cortical excitatory neurons^33^. Specifically, we used monoclonal KOLF2.1J-CRISPRi-Ngn2 cell lines carrying a constitutively expressed CRISPRi cassette in the CLYBL safe harbor locus as well as a Neurogenin induction cassette in the AAVS1 locus for inducing a cortical neuron-like state. These human induced pluripotent stem cells (iPSCs) take approximately 7 days to differentiate into cortical-like neurons (i3Neurons)^34,35^ following established protocols^36^ and as depicted in **Fig 1A**. Please see the Methods section for additional details on cell culture and differentiation.

**Figure 1:**
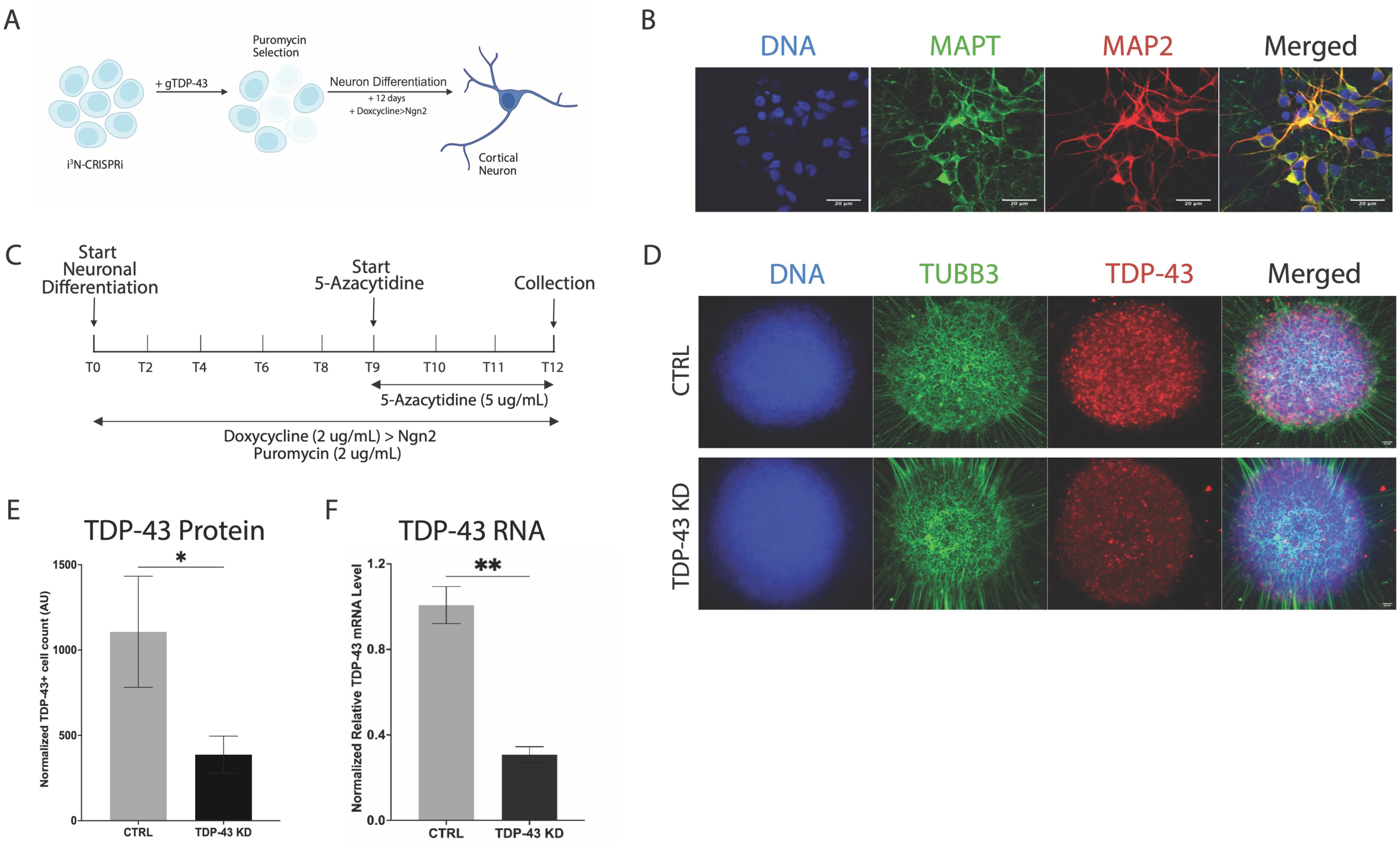
Generation of neuronal-like models of TDP-43 nuclear loss. (**A**) Schematic of the i3Neuron differentiation protocol from CRISPRi-i3N-KOLF2.J1 iPSC lines. (**B**) Immunofluorescence of cortical excitatory neuron markers expressed by mature i3Neurons include MAPT (green), MAP2 (red), and the DNA marker DAPI (blue). Images were obtained at 40x magnification with scale bars indicating a 20μM distance. (**C**) A timeline of i3Neuron differentiation following guide RNA (gRNA) transduction for TDP-43 or non-targeting control gRNAs. Time for addition of 5-Azacytidine is also shown. (**D**) Immunofluorescence detection of TDP-43 protein (red) in i3Neuron cells transduced with control (top) or TDP-43 targeting gRNAs (bottom). The neuronal specific marker TUBB3 (green) as well as DAPI (blue) remain unchanged in TDP-43 KD cells. **(E)** Quantification of immunofluorescence identification of TDP-43 positive i3Neuron cells transduced with control or TDP-43 targeting gRNAs (mean ± SEM, N=12 CTRL, N=15 TDP-43 KD, P<0.0308, two-tailed t-test). **(F)** Relative TDP-43 mRNA level calculated by normalization of TDP-43 mRNA level by RPL13A using qPCR for i3Neurons expressing the TDP-43 targeting gRNA vs the non-targeting gRNA (mean ± SEM, three biological replicates) (log2FC = –1.74, P<0.0018, two-tailed t-test).

We verified that these i3Neurons express genes typical of cortical excitatory neurons at day 15 post-differentiation using immunofluorescence staining for the neuronal markers MAPT (green) and MAP2 (red) (**Fig 1B**). We also included in our model the addition of 5-Azacytidine, 5-Aza, which has previously been reported to inhibit non-sense mediated decay (NMD) of RNA transcripts^37^ in order to allow for enhanced capture of cryptic splice isoforms which would otherwise be degraded by NMD (**Fig 1C**). These cells were transduced with two guide RNAs (gRNAs) targeting the TARDBP locus to achieve at least 2-fold loss of TDP-43 expression at the RNA and protein level. To confirm TDP-43 knockdown in our i3Neurons, immunofluorescence staining was carried out for TDP-43 (red), the neuronal marker TUBB3 (green) and DAPI (blue) (**Fig 1D**). Quantification of the number of TDP-43 positive cells showed an approximate 3-fold reduction (p-value < 0.031, two-tailed t-test) following TDP-43 knockdown (TDP-43 KD) (**Fig 1E**). Changes in TARDBP RNA levels were assessed with qPCR and also showed substantial and significant RNA reduction (log_2_FC = –1.74, p-value < 0.002, two-tailed t-test) (**Fig 1F**).

### TDP-43 depletion in i3Neurons results in dysregulation of known TDP-43 splice targets and elevation of transposable elements (TEs)

Expression and splicing profiles of our i3Neurons were generated from samples collected on day 12 post-differentiation (**Fig 2A**). Short-read RNA sequencing (RNA-seq) libraries were constructed for two replicates of each of the 4 conditions (**Fig 2B**): i3Neurons transduced with two non-targeting gRNAs (CTRL), i3Neurons transduced with non-targeting gRNAs and 5-Azacytidine treatment (5-Aza), i3Neurons transduced with two gRNAs targeting TDP-43 (TDP-43 KD), and i3Neurons transduced with two gRNAs targeting TDP-43 and treated with 5-Azacytidine (TDP-43+5-Aza). A principal components analysis (PCA) plot based on gene expression profiles from all samples (**Fig 2B**) shows clear clustering by TDP-43 KD/control and 5-Aza treatment status. Differential gene expression analysis returned 10,329 significantly altered genes in the TDP-43+5Aza samples, as compared to controls. A volcano plot in **Fig 2C** highlights known TDP-43 target genes^38^ (MALAT1^39,40^, STMN2^9^, PTPRN2^41^, HDGFL2^41^) as well as the gene encoding TDP-43 itself (TARDBP)^42^. All differentially expressed genes are listed in supplemental tables **S1A-C**.

**Figure 2:**
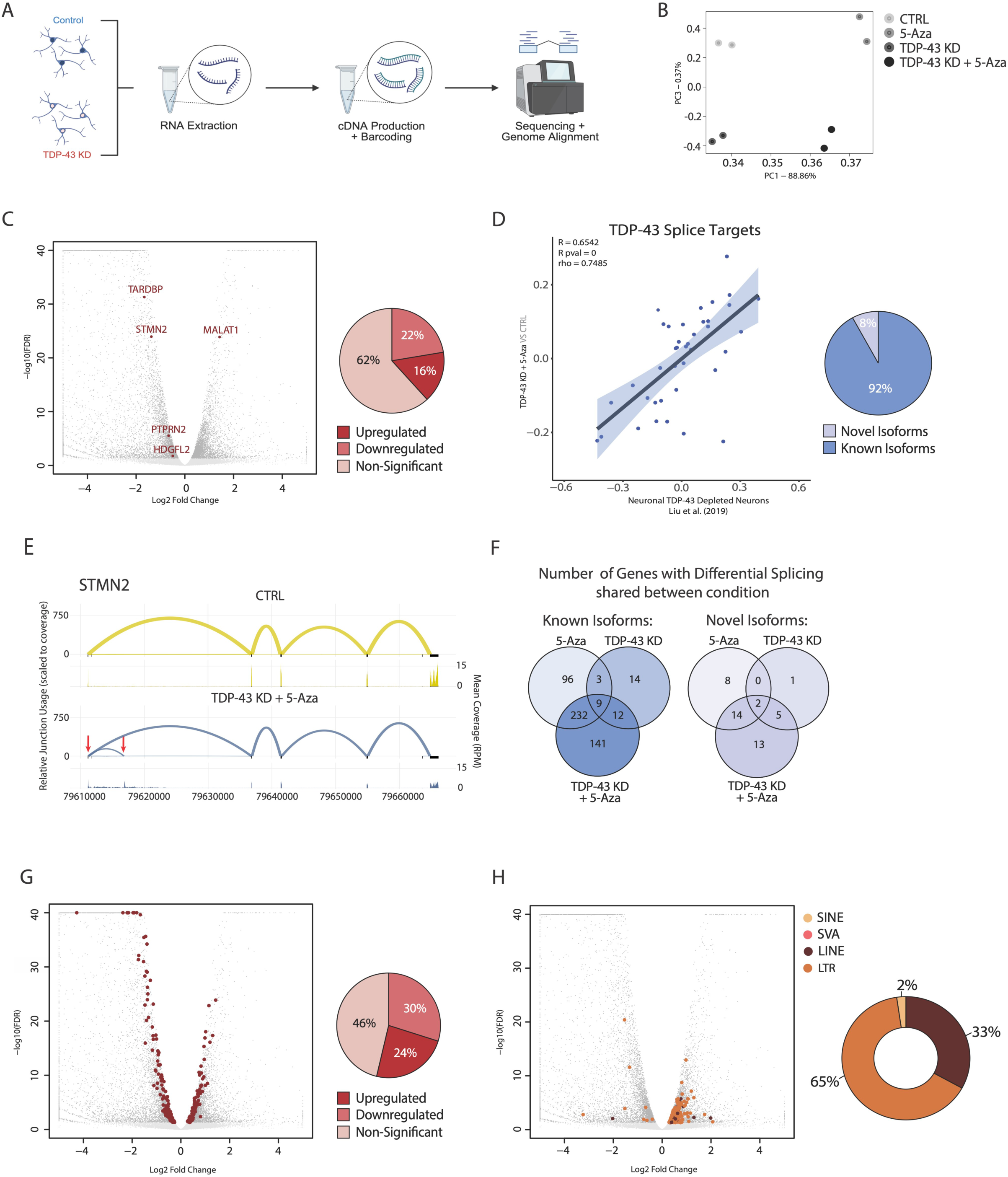
Detection of known and novel splice isoforms in TDP-43 depleted i3Neurons. **(A)** Schematic of RNA-Seq workflow for collection and sequencing of i3Neurons which received non-targeting or TDP-43 targeting gRNAs. **(B)** PCA i3Neuron RNA-seq libraries for samples transduced with either non-targeting (CTRL) or TDP-43 targeting (TDP-43 KD) gRNAs and with/without 5-Azacytidine (5-Aza) treatment **(C)** Volcano plot of fold change vs false discovery rate (FDR) q values for genes from i3Neurons with TDP-43 targeting gRNAs and 5-Azacytidine treatment. Significantly differentially expressed genes (dark grey) and known TDP-43 regulated gene targets (red) are highlighted. The pie chart shows percentages of significantly upregulated and downregulated genes. **(D)** Scatter plot of genes with splicing changes (PSI (Ψ)) detected in sorted ALS/FTD neurons depleted for TDP-43 vs TDP-43+5-Aza i3Neurons (Pearson’s R=0.65, P<1E-10). Pie chart of known and novel isoforms detected in TDP-43+5-Aza i3Neurons. **(E)** Sashimi plot of relative junction usage scaled to coverage (top) and corresponding mean coverage (RPM) (bottom) from Leafcutter analysis of STMN2 from CTRL (yellow) and TDP-43+5-Aza (blue) i3Neurons (two biological replicates). Red arrows indicate cryptic exon inclusion events specific to TDP-43 loss. (**F)** Venn diagrams showing genes with significant differential splicing, creating known and novel isoforms, compared to CTRL which are shared between all treatments in i3Neurons. **(G)** Volcano plot of fold change vs false discovery rate (FDR) q values for genes from TDP-43+5-Aza i3Neurons. Genes with both significant differential expression and differential splicing compared to CTRL are highlighted (red). The pie chart displays the percentage of differentially spliced genes that are up– or down-regulated. **(H)** Volcano plot of fold change vs false discovery rate (FDR) q values for significantly differentially expressed genes (dark grey) and TE subfamilies (SINE = gold, SVA = pink, LINE = maroon, LTR = orange) in TDP-43+5-Aza i3Neurons. The donut plot shows the percentage by family of these differentially expressed TE subfamilies that show upregulation.

We next assessed TDP-43 dependent changes in splice junction usage via LeafCutter^43^, which returned 1617 significantly altered splice sites in the TDP-43+5-Aza samples as compared to controls. These altered splice site usage measurements are expressed as changes in the percent spliced in (PSI (Ψ)) of a particular exon junction. Changes in PSI include both canonical splice sites as well as cryptic (novel) splice sites, not previously annotated as part of a gene isoform (see supplemental table **S2**A). To verify the extent to which our i3Neuron libraries recapitulate the cryptic splicing patterns observed in ALS/FTD patient tissue, we compared the splicing dysfunction observed in our TDP-43+5-Aza i3Neurons with a previously published data set of sorted TDP-43 negative neurons from ALS/FTD post-mortem brain samples, as profiled by Liu et al^29^. We found a significant correlation (Pearson’s R=0.65, p-value < 1.0^−10^, **Fig 2D**), between splicing changes (PSI (Ψ)) in our TDP-43 knockdown i3Neurons and the sorted ALS/FTD neurons that were depleted for TDP-43, indicating that our model for TDP-43 neuronal loss of function recapitulated a significant proportion of TDP-43 dependent cryptic splicing defects found in ALS/FTD patients. Of the total detected differential splicing events in our TDP-43+5-Aza vs control i3Neurons, the majority (92%) represented known splice junctions as annotated in GCv47 (see methods), with the remaining 8% representing novel (cryptic) splicing events not annotated in GCv47 (**Fig 2D**, Supplemental Table **S3**). An example sashimi plot highlighting one previously reported TDP-43 dependent cryptic splicing event (STMN2^9,13^) is shown in **Fig 2E**, where the canonical STMN2 splice pattern is shown for controls (yellow), the detected STMN2 splice patterns in our TDP-43+5-Aza i3Neurons is shown in blue, and the cryptic splice junctions are highlighted by red arrows.

Comparing genes with differential splicing events across treatments in i3Neurons, TDP-43+5Aza accounts for the largest number of genes with known and novel splice isoforms (**Fig 2F**). Next, we evaluated whether TDP-43 dependent cryptic splicing was also associated with differential gene expression of those TDP-43 splice targets. Of the genes with detected TDP-43 mediated cryptic splicing events, 24% showed significant upregulation and 30% displayed significant downregulation (**Fig 2G**). This indicates that a majority of TDP-43 dependent cryptic splice alterations are accompanied by gene expression changes, and that these changes involve both increases and decreases in gene expression.

Finally, we also assessed whether our i3Neurons show TDP-43 dependent changes in the expression of transposable elements (TEs). Previous work from our group^20^ and others^30^ have shown that TDP-43 dysfunction leads to elevation in the expression of broad categories of human TEs. This includes several subfamilies of long-terminal repeat retrotransposons (LTRs), long interspersed nuclear elements (LINEs), short interspersed nuclear elements (SINEs), and the human specific SINE-VNTR-Alu (SVA) elements. Consistent with previous studies of TDP-43 dependent TE de-silencing^20,30^, we saw broad upregulation of genomic repeats from every major TE family in the i3Neurons upon TDP-43 loss and 5-Aza treatment (**Fig 2H**). LTRs made up nearly two thirds of the elevated TEs (65%), followed by LINEs contributing at 33%, and with the remaining 2% attributed to the SINE subfamily (**Fig 2H**, volcano plot and pie chart). In summary, the i3Neuron models recapitulate splicing alterations and gene/TE expression changes previously seen in cellular models of TDP-43 loss^9,20,41^, ALS-FTD cortical and spinal cord tissues^13,44,45^, as well as sorted ALS/FTD patient neurons^29^.

### Long read RNA sequencing of TDP-43 depleted neurons reveals thousands of novel gene transcripts and full-length transposable elements (TEs)

To evaluate the contributions of TDP-43 neuronal loss to cryptic splicing and dysregulation of TE transcripts, we required a sequencing technique to allow for the capture and resolution of full-length RNA transcripts. Short read RNA-seq profiles allow for highly quantitative measures of gene expression and exon junction inclusion rates, but are less optimal for full RNA transcript assembly and for resolving repetitive sequences, such as TEs^46,47^. To overcome these limitations, we developed a modified method for long read RNA sequencing (**Fig 3A**), based upon the PacBio Iso-seq protocols, which uses a highly processive reverse transcriptase enzyme (MarathonRT)^48^ to optimize reverse transcription of long and highly structured transcripts (please see Methods for further details). This method allowed for the efficient capture of full-length gene mRNAs as well as full-length TE transcripts. In particular, the low error rates of the PacBio Iso-seq platform enabled the resolution of TE-derived transcripts to the specific originating TE locus. Furthermore, this method is optimized for both gene and TE transcript sequencing, such that the majority of our reads represent known isoforms of annotated genes (86%, **Fig 3B**), while also recovering thousands of novel isoforms and novel genes across all conditions (**Fig 3B** and supplemental tables **S4A-B**).

**Figure 3:**
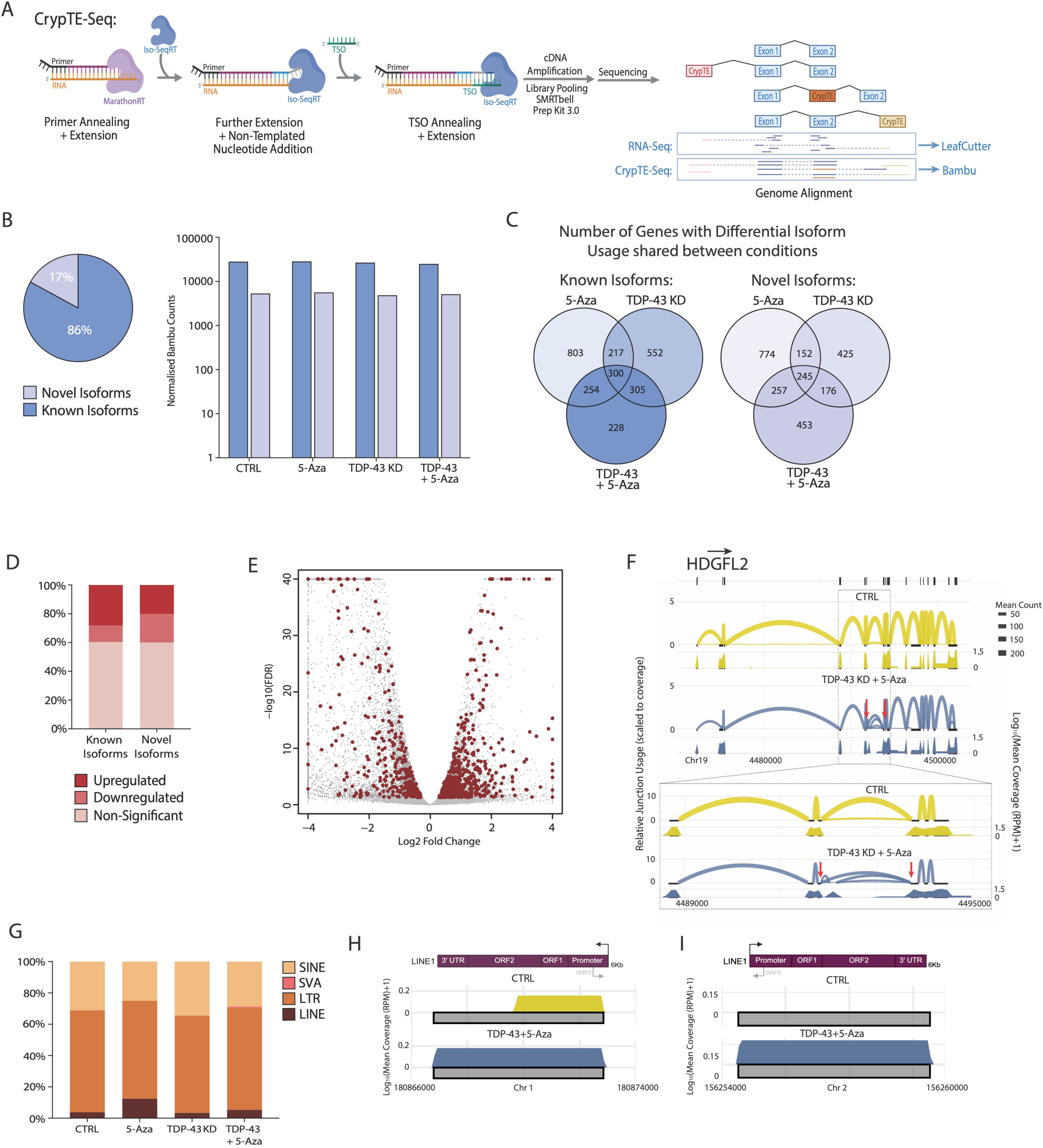
Long-read CrypTE-seq libraries improve the detection of cryptic splice isoforms and reveal thousands of novel crypTE events in TDP-43 depleted i3Neurons. **(A)** Schematic of the CrypTE-Seq method highlighting the introduction of MarathonRT prior to continuation with the standard Iso-Seq protocol and genome alignment of sequencing reads. **(B)** Pie chart of the percentage of known (blue) and novel (lilac) isoforms detected with Bambu analysis of CrypTE-Seq libraries in TDP-43+5-Aza i3Neurons. Grouped bar charts display for each i3Neuron condition the number of normalized counts per isoform type (known = blue, novel = lilac). **(C)** Venn diagrams showing genes with differential isoform usage, creating known and novel isoforms, compared to CTRL which are shared between all treatments in i3Neurons. **(D)** Stacked bar charts for known and novel isoforms detected in TDP-43+5-Aza i3Neurons with CrypTE-Seq and the percentage that are significantly up-/down-regulated compared to CTRL. **(E)** Volcano plot of fold change vs false discovery rate (FDR) q values for significantly differentially expressed genes (dark grey) from matched RNA-Seq data for TDP-43+5-Aza i3Neurons highlighting (red) the genes with CrypTE-Seq identified differential isoform usage. **(F)** Sashimi plot of relative junction usage in the HDGFL2 gene scaled to coverage (top) and corresponding binned mean coverage (RPM+1) (bottom) for CTRL (yellow) and TDP-43+5-Aza (blue) i3Neurons CrypTE-Seq libraries. Red arrows indicate a known TDP-43 exon inclusion event which occurs with TDP-43 loss of function. **(G)** Stacked bar chart showing the proportions of each TE subfamily (DNA = green, SINE = gold, SVA = pink, LINE = maroon, LTR = orange) detected with CrypTE-Seq per i3Neuron condition. (**H-I**) Coverage (log_10_ mean coverage (RPM+1)) of CrypTE-seq reads over intact (H) L1HS and (I) L1PA2 LINE elements showing that reads begin and end at the annotated LINE-1 boundaries in TDP-43+5-Aza i3Neurons.

Focusing on the differentially spliced genes, we detected over 5 times more genes with differential isoform usage events per i3Neuron condition in our long-read libraries (**Fig 3C**, supplemental table **S4B**) as compared to the short-read based data (**Fig 2F**). A substantial fraction of the novel isoforms detected in our study were also associated with differential expression at the gene level, with 40% of novel isoforms showing either significant elevation (20%) or depletion (20%) at the gene level in TDP43+5Aza as compared to controls (**Fig 3D**). These were comparable to gene expression changes associated with annotated isoforms, where 28% showed upregulation and 12% were significantly downregulated (**Fig 3D**). Figure **3E** displays a volcano plot of differential gene expression in TDP-43+5-Aza i3Neurons vs. controls, where genes that were both differentially spliced in the CrypTE-seq TDP-43+5-Aza vs control libraries and differentially expressed at the RNA-Seq level are marked in red.

The isoforms with differential usage detected in our TDP-43+5-Aza long read libraries included a majority of the previously published TDP-43 dependent cryptic splice isoforms. An example sashimi plot for HDGFL2 is shown in **Fig 3F**. HDGFL2 contains a TDP-43 dependent cryptic exon inclusion event previously reported by Seddighi et al.^41^ and shows evidence for the same TDP-43 cryptic splicing event in our TDP-43+5-Aza i3Neurons. Overall, we detected the same cryptic splice junctions that define TDP-43 cryptic isoforms for 74% of the transcripts reported in Ma et al^12^ (51/69=74%) and 63% of the transcripts reported in Seddighi et al^41^ (279/445=63%). For the full list of overlapping cryptic splice junctions across studies, please see Supplemental Table **S5**.

We next explored the ability of the CrypTE-seq assays to return expressed TE transcripts. For the TE transcripts that were detected in our long-read libraries, we first summed the read counts for each subfamily per condition and calculated the percentage of reads each subfamily contributes to the overall number of TE transcripts for that treatment (**Fig 3G**). HERV elements are poorly captured by long read sequencing techniques that employ polyA capture steps, such as Iso-Seq, as HERVs are thought to produce RNA transcripts processed more similarly to long non-coding RNAs which show reduced polyadenylation^49^. For this reason, we’ve highlighted 5’ intact LTR elements which can exist as solo LTRs or act as promoters to drive TE or gene expression. Due to this annotation bias, LTR driven sequences make up a large percentage of the TE transcripts that were captured by CrypTE-Seq across all i3Neuron conditions (CTRL 65%, 5-Aza 63%, TDP-43 KD 62%, TDP-43+5-Aza 65%). The next most abundant TE subfamily with transcripts captured are SINEs (CTRL 31%, 5-Aza 25%, TDP-43 KD 34%, TDP-43+5-Aza 29%). This is consistent with their abundance, representing approximately 11% of the human genome^50^.

We next took advantage of the locus-specific resolution of long-read libraries to identify individual TEs that are independently transcribed from their own promoters. Many TE copies in the genome represent fragments due to recombination, structural variation, and truncation during aborted transposition attempts^51,52^. In particular, many TEs are missing their 5’ sequences and do not contain intact promoters. Thus, we filtered the human annotated TE list to select those that contain intact 5’ ends and used this list to search for intact TE transcripts. We found hundreds of intact TE transcripts captured in each condition (supplemental table **S6A-D**) including many human-specific and young TEs. Examples of full length and intact LINE-1 elements captured by CrypTE-Seq in i3Neurons with TDP-43+5-Aza treatment are highlighted in **Fig 3H-I**. Both of these LINE-1 elements (L1HS: **Fig 3H** and L1PA2: **Fig 3I**) span the full 6kb length of an intact LINE, contain no truncating mutations, and have previously been captured in lower-throughput RACE assays targeting expressed LINE-1 sequences^53^. L1HS elements are particularly intriguing because they encode all the proteins necessary to produce cDNA, proteins, and to autonomously transpose^54^. Of note, the L1HS element shown in **Fig 3H** derives from an intronic location (an intron of XPR1), which was true of many of the TE transcripts captured. Yet, the long read capture strategy enabled us to determine that the reads began and ended at the annotated ends of each TE locus and did not derive from read-through transcription of overlapping or adjacent genes, as displayed in the read coverage plots (**Fig 3H**). This full-length 6kb L1PA2 element (**Fig 3I**) has been previously reported to be transcriptionally active in human neuronal precursors^55^, where activation of its alternative antisense promoter (ORF0) overlaps the start of an adjacent long non-coding (lncRNA) LINC01876 and drives expression of this lncRNA during human brain development.

### Long read CrypTE-seq assays uncover widespread cryptic splicing between genes and adjacent TEs that accumulate in TDP-43 depleted neurons

Due to the dual role of TDP-43 in the regulation of TE expression and RNA splicing fidelity, we next asked whether some of the TE sequences in our libraries might derive from cryptic splicing between genes and TEs, or crypTEs. Depending on the location of the TE relative to a gene, there can be different consequences for cryptic splicing events. For example, upstream TEs can splice into downstream genes and act as alternative transcription start sites (crypTE-TSS, **Fig 4A**). Intronic TEs can be included as cassette exons within gene transcripts (crypTE-Exon, **Fig 4B**). Upstream genes can splice into and terminate transcription within downstream TEs, using the TE as an alternate polyadenylation site (crypTE-ApA, **Fig 4C**). To distinguish these cryptic splicing events from cases of potential transcriptional read-through or simple intron retention, we chose to restrict the list of crypTE isoforms to those events which used a splice donor and/or acceptor site within the TE itself. Given that the majority of the TE-annotated reads in our long-read libraries were derived from these crypTE transcripts, we named our sequencing assay CrypTE-seq. In all, the CrypTE-Seq assays enabled the capture of full-length gene mRNA isoforms, cryptic gene isoforms that did not involve TEs, full-length TE transcripts, and cryptic gene-TE splice fusions (**Fig 4D**). As shown in **Fig 4D**, the majority of the cryptic transcripts detected involved either non-TE cryptic splice events (such as those present in STMN2 or HDGFL2) or crypTE-Exon inclusions in TDP-43+5-Aza i3Neurons.

**Figure 4:**
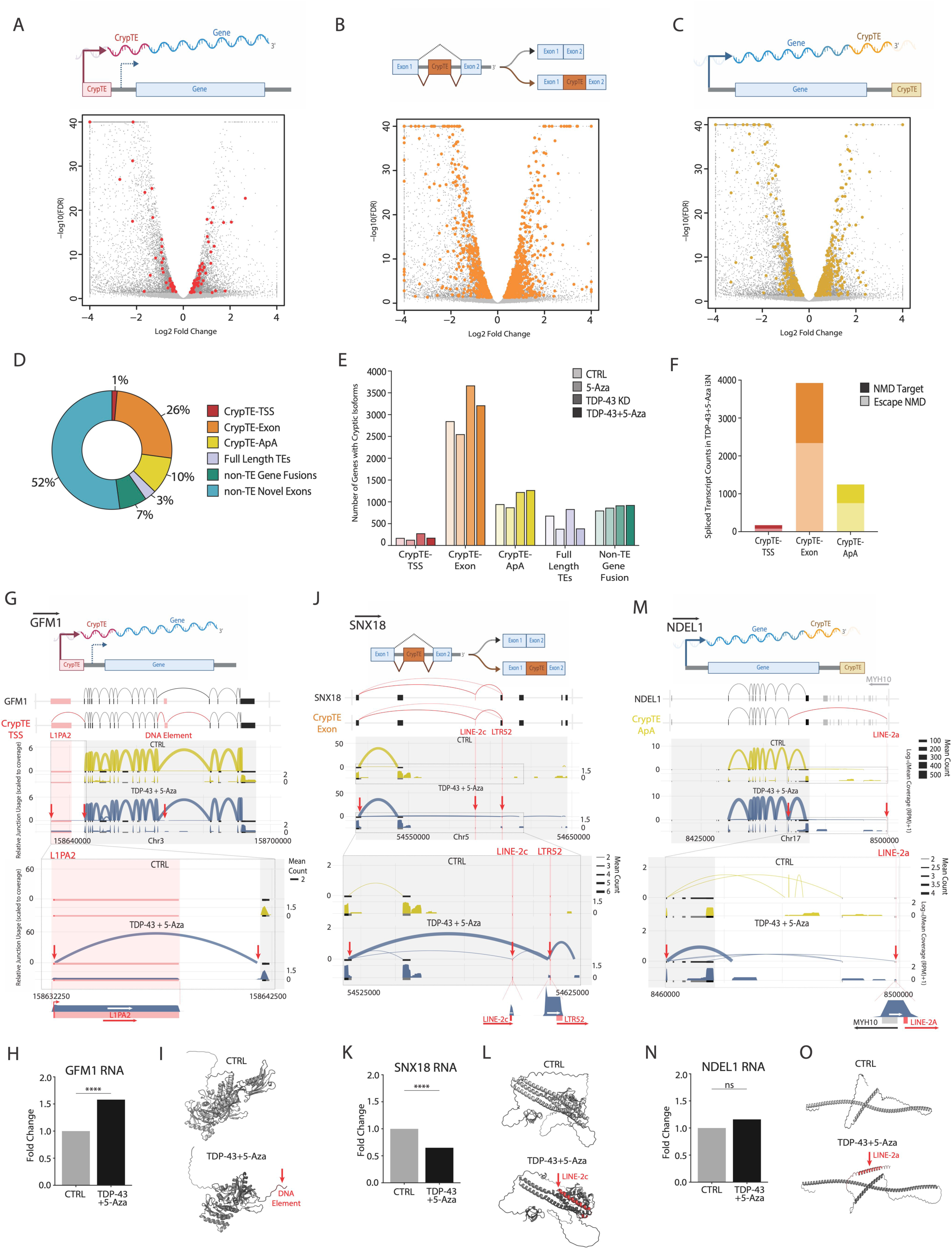
CrypTE transcripts are associated with changes in gene expression and are predicted to produce novel proteins from TE sequences. **(A-C)** Schematic of **(A)** crypTE-TSS, **(B)** crypTE-Exon and **(C)** crypTE-ApA events. Below each schematic are volcano plots of fold change vs false discovery rate (FDR) q values for significantly differentially expressed genes (dark grey) from matched RNA-Seq data for TDP-43+5-Aza i3Neurons highlighting genes with **(A)** crypTE-TSS (red), **(B)** crypTE-Exon (orange) or **(C)** crypTE-ApA (yellow) events detected by CrypTE-Seq. **(D)** A donut chart displays the proportion for each category of cryptic splicing event captured by CrypTE-Seq (non-TE Novel Exons = blue, CrypTE-TSS = red, CrypTE-Exon = orange, CrypTE-ApA = yellow, Full-length TEs = purple, non-TE Gene Fusions = green) in TDP-43+5-Aza i3Neurons. **(E)** Grouped bar charts show the number of genes detected with CrypTE-Seq containing each crypTE event type (crypTE-TSS = red, crypTE-Exon = orange, crypTE-ApA = yellow), full-length TEs (purple) and non-TE gene fusion events (green) by treatment type of i3Neurons (shading indicates i3Neuron condition from lightest to darkest: CTRL, 5-Aza, TDP-43 KD, TDP-43+5-Aza). **(F)** Stacked bar charts for the number of spliced transcripts that are predicted to alert non-sense mediated decay (NMD) pathways (top, dark shading) or to escape NMD (bottom, light shading) in TDP-43+5-Aza i3Neurons. **(G, J, I)** Schematics highlighting the splicing patterns (black boxes = gene exons, pink boxes = TEs) of a canonical splice isoform (top) and the crypTE containing isoform (below) for the genes: **(G)** GFM1, **(J)** SNX18 and **(M)** NDEL1. Directly below these are the sashimi plots of relative junction usage scaled to coverage (top) and corresponding binned mean coverage (log_10_(RPM+1)) (bottom) from CTRL (yellow) and TDP-43+5-Aza (blue) i3Neurons CrypTE-Seq libraries for **(G)** GFM1, **(H)** SNX18 and **(I)** NDEL1. Red arrows indicate crypTE splicing events found in TDP-43+5-Aza but not CTRL i3Neurons **(H, K, N)** Bar charts of fold change RNA expression in TDP43+5-Aza i3Neurons normalized to control expression from RNA-Seq data for each of the crypTE containing genes highlighted: **(H)** GFM1 (1.58 fold upregulation, P_adj_ < 3.159×10^−5^), **(K)** SNX18 (1.53 fold down-regulation, P_adj_ < 2.138×10^−6^) and **(N)** NDEL1 (1.16 fold upregulation, P_adj_ < 0.186). **(I, L, O)** AlphaFold 3 protein structure predictions of **(I)** GFM1, **(L)** SNX18 and **(O)** NDEL1 canonical transcripts in CTRL (top, light grey) and crypTE containing transcripts in TDP-43+5-Aza (bottom, dark grey = gene, red = TE).

Many of these crypTE transcripts were associated with significant differential expression of the associated gene in the TDP-43+5-Aza i3Neurons. For example, crypTE transcripts driven from TE promoters tend to result in increased total expression of the associated gene, as shown in the volcano plot of **Fig 4A**. In this volcano plot, Log2 fold change in gene expression in the TDP-43+5-Aza libraries compared to controls is shown on the horizontal axis, Log10 adjusted P-values (FDR) is plotted on the vertical axis, all differentially expressed genes are colored dark grey, and genes associated with crypTE-TSS events are colored red (**Fig 4A**, bottom). This indicates that adding an additional TE promoter tends to be additive rather than interfering with normal gene transcription from the gene TSS. Volcano plots are also shown for crypTE events resulting from cassette TE exon inclusion (crypTE-Exon, orange dots, **Fig 4B**) and alternative 3’ UTR/polyadenylation events (crypTE-ApA, yellow dots, **Fig 4C**), where we noted both up– and down-regulation of the associated genes in the TDP-43+5-Aza libraries.

There were clear differences in the number of crypTE events captured. As shown in the pie charts of **Fig 4D** and the bar plots of **Fig 4E**, crypTE-exon events were the most common type in our libraries, followed by crypTE-ApA events, both of which showed increases in response to TDP-43 KD (**Fig 4E**, supplemental table **S7**). Due to the technical limitations associated with long-read sequencing techniques (3’ polyA capture bias), the number of true crypTE-TSS events may be higher than the transcripts we detected (1%), which were the least common type of crypTE transcript. Full length TEs were restricted to elements that began at the TE promoter, without evidence for read-through and were less common than crypTE transcripts (3%). The remainder of the novel isoforms in TDP-43_5-Aza i3Neurons detected by CrypTE-seq did not involve TEs (supplemental tables **S8-9**). These non-TE events included both cryptic splice isoforms (52%) and gene-gene fusion events (7%). Looking across conditions for i3Neurons, each of these novel isoform types and full-length TE transcripts were detected at some level in control, untreated cells (**Fig 4E**). We detected a fair number of crypTE events even in unperturbed i3Neurons (**Fig 4E**), indicating that TE exonization occurs at basal rates in normal cells. Importantly, the production and/or accumulation of thousands of crypTE transcripts increases in the absence of TDP-43 (supplemental table **S7**).

### CrypTE isoforms may impact gene function through creating premature termination codons or through production of novel peptides

CrypTE containing sequences within gene transcripts can lead to novel or detrimental impacts on overall gene function. When crypTE sequences provide premature termination codons, this can alter total gene levels through promoting nonsense mediated decay (NMD) of the transcript^56,57^. When crypTE sequences occur within the open reading frame (ORF) of this transcript, this can result in the production of a novel peptide that translates portions of the crypTE sequence with potential downstream consequences on protein function. To address this, we assessed the fraction of crypTE transcripts that contain premature termination codons and thus are predicted to be NMD targets (**Fig 4F**). For each type of crypTE transcript in the TDP-43+5-Aza i3Neurons, we assessed whether the crypTE sequence was likely to occur within a potential ORF and, if so, whether this would generate premature termination codons that could alert NMD machinery (see supplemental table **S10**)^58^. As shown in **Fig 4F**, there were similar proportions of transcripts of each crypTE type that were predicted to be NMD targets. Specifically, 60% of both the crypTE-Exon and crypTE-ApA transcripts were predicted NMD targets. A smaller fraction of the crypTE-TSS transcripts were predicted NMD targets (45%), which may be consistent with the fact that these transcripts were also less likely to be down-regulated at the expression level.

For each crypTE event type, we highlighted a specific example that demonstrates these impacts on gene expression and potential for peptide formation. One example crypTE-TSS event involved the use of a LINE-1 (L1PA2) promoter to drive expression of its immediate downstream neighbor, GFM1 (**Fig 4G**). In this case, the TDP-43KD+5Aza libraries showed both expression of the full-length 6kb L1PA2 element itself as well as splicing from the LINE-1 promoter into the 5’UTR region of GFM1, neither of which were detected in control libraries. Using the highly quantitative matched short-read RNA-seq data, we determined that total GFM1 gene expression is increased in TDP-43KD+5Aza libraries as compared to controls (1.58 fold upregulation, P_adj_ < 3.159×10^−5^, **Fig 4H**). Furthermore, this particular LINE-1 driven GFM1 crypTE transcript also terminates within another downstream TE sequence (a Tigger element), suggesting that the resulting transcript would not code for its canonical GFM1 protein. We used AlphaFold^59^ to determine whether this crypTE transcript is likely to produce a novel protein (see Supplemental table **S11** for a full list of AlphaFold predicted peptides). AlphaFold predicts a truncated protein would be produced, with novel TE sequence included in the C-terminal domain of GFM1-cryptic (**Fig 4I**). GFM1 is a mitochondrial elongation factor and is necessary for the oxidative phosphorylation process^60,61^. Despite the expression of GFM1 in all cell types, mutations in GFM1 typically induce neurological symptoms^62,63^.

An example crypTE-exon transcript involved two short fragmented TEs (LINE2c and LTR-52), which are included as part of cryptic exons within the sorting nexin gene, SNX18. SNX18 plays roles in endocytosis^64–66^ and autophagosome formation^67,68^. The SNX18 crypTE-exon transcript (**Fig 4J**) skips exon 2 of SNX18 and instead splices into two adjacent TEs (LINE2c and LTR-52), before terminating downstream. This event is also associated with significant down-regulation of total SNX18 gene levels (1.53 fold down-regulation, P_adj_ < 2.138×10^−6^, **Fig 4K**). AlphaFold predicts inclusion of LINE-2c sequence as part of a novel domain near the C-terminus of the cryptic SNX18 protein (**Fig 4L)**.

Finally, an example crypTE-ApA transcript fuses the NDEL1 gene transcript to a LINE-2A sequence that lies approximately 30kb downstream of NDEL1 (**Fig 4M**). This LINE-2A termination sequence replaces the last canonical exon (exon 11) of NDEL1 with cryptic TE-containing sequence. In this case, we did not detect significant differential expression of total NDEL1 RNA levels in TDP-43+5-Aza i3Neurons (**Fig 4N**). However, the LINE-2A sequence is predicted by AlphaFold to form a new protein domain within the C-terminal region as part of a novel fusion peptide (**Fig 4O**). NDEL1 is known to play roles in dynein complex mediated transport^69–71^ and neurofilament polarization^72^, where the C-terminal domain is required for proper protein-protein interactions^73,74^.

### CrypTE isoforms accumulate in ALS/FTD frontal cortex tissues with TDP-43 pathology and cognitive impairment

We next explored the extent to which these crypTE transcripts are detectable in ALS/FTD frontal cortex tissues and associated with TDP-43 pathology and brain region-relevant clinical parameters, such as cognitive impairment. To that end, we generated CrypTE-Seq libraries from 8 post-mortem frontal cortex (BA46) samples from ALS/FTD patients that had both frontal cortex TDP-43 pathology and clinical signs of cognitive impairment (ALS-CI, n=3) or from ALS/FTD patients that did not present with cognitive impairment or TDP-43 pathology (ALS-nonCI, n=3) as well as two non-diseased controls (CTRL, n=2) as displayed in **Fig 5A**. Specifically, these tissues were from the BA46 region of frontal cortex, which has been associated with executive function and working memory^75^. Quantitative measures of cognitive impairment and detection of TDP-43 pathology are available for all subjects as described previously^75^ (supplemental table **S12**). We first verified that these CrypTE-seq libraries were of similar quality to our i3Neuron libraries, with similar levels of total detected transcripts (supplemental tables **S4A, S13A**), similar fractions of known and novel genes detected (supplemental tables **S4B, S13B**), and presence of previously published TDP-43 cryptic splicing events^12,41^ (supplemental table **S13C**).

**Figure 5:**
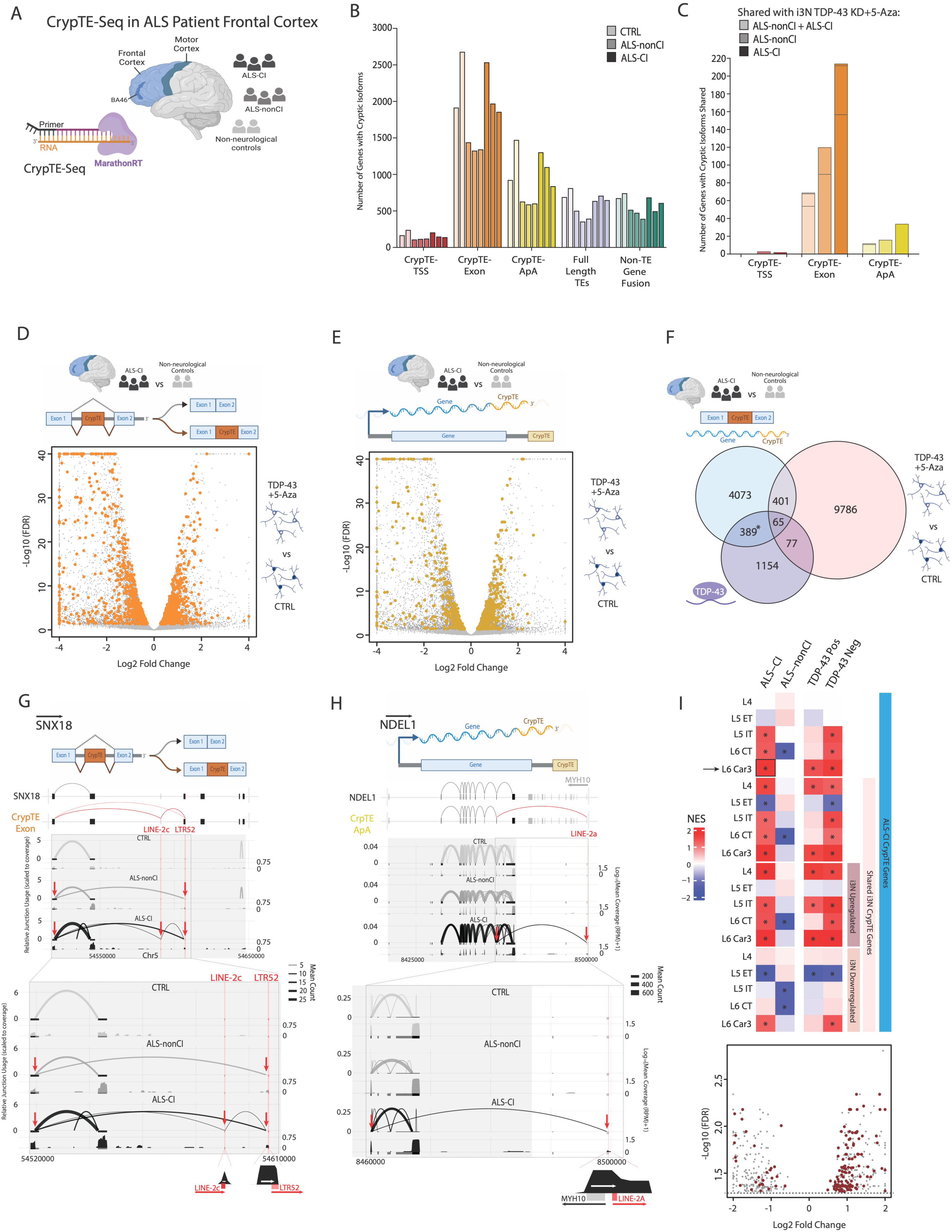
TDP-43 mediated crypTE transcripts are detected in ALS/FTD frontal cortex tissues and associated with TDP-43 dysfunction. **(A)** Schematics for CrypTE-Seq library generation from post-mortem frontal cortex tissues from Brodmann area 46 (BA46) in ALS patients diagnosed with cognitive impairment (ALS-CI) and without cognitive impairment (ALS-nonCI). Control (CTRL) tissues were obtained from non-neurological age-matched individuals. **(B)** Grouped bar charts showing the number of genes detected using CrypTE-Seq which contained each crypTE event type (crypTE-TSS = red, crypTE-Exon = orange, crypTE-ApA = yellow), full-length TEs (purple) and non-TE gene fusion events (green) identified for each patient (shading indicates group from lightest to darkest: CTRL, ALS-nonCI, ALS-CI). **(C)** Grouped bar chart for each crypTE category (CrypTE-TSS = red, CrypTE-Exon = orange, CrypTE-ApA = yellow) showing the number of genes detected with CrypTE-Seq containing the exact same crypTE events in TDP-43+5-Aza vs CTRL i3Neurons and ALS-nonCI and/or ALS-CI patients which are absent in non-neurological controls (shading from lightest to darkest indicates sharing between TDP-43+5-Aza i3Neurons and both ALS-CI and ALS-nonCI, ALS-nonCI only, ALS-CI only). Stacked crypTE-Exon bars include crypTE-Exon only genes (bottom), crypTE-Exon/ApA (middle) and crypTE-Exon/TSS genes (top). Stacked crypTE-ApA bars include crypTE-ApA (bottom) and crypTE-ApA/TSS (top) genes. **(D-E)** Volcano plots of fold change vs false discovery rate (FDR) q values for significantly differentially expressed genes (dark grey) from RNA-Seq data for TDP-43+5-Aza i3Neurons highlighting genes from ALS-CI patients with **(D)** crypTE-Exon (orange) or **(E)** crypTE-ApA (yellow) events not in non-neurological controls detected by CrypTE-Seq. **(F)** Venn diagram comparing genes which are differentially expressed in TDP-43+5-Aza vs control i3Neurons (pink), eCLIP targets of TDP-43 protein in SH-SY5Y (purple) and contain crypTE-Exon and/or crypTE-ApA events in ALS-CI patient BA46 tissue not in non-neurological controls. **(G-H)** Schematics highlighting the splicing patterns (black boxes = gene exons, pink boxes = TEs) of a canonical splice isoform (top) and the crypTE containing isoform (below) for the genes: **(G)** SNX18 and **(H)** NDEL1. Directly below these are the sashimi plots of relative junction usage scaled to coverage (top) and corresponding binned mean coverage (log_10_(RPM+1)) (bottom) from CTRL (light grey), ALS-nonCI (dark grey) and ALS-CI (black) patient post-mortem frontal cortex from BA46. Red arrows indicate crypTE splicing events found in ALS-nonCI (n=3) and/or ALS-CI (n=3) but absent in CTRLs (n=2). **(I)** Heatmap (top panel) with each row representing the differentially expressed genes from snRNA-Seq in the excitatory neurons corresponding from that BA46 grey matter layer (layer 4 = L4, layer 5 extratelencephalic (ET) = L5 ET, L5 intratelencephalic (IT) = L5 IT, layer 6 corticothalamic (CT) = L6 CT and L6 Car3 intratelencephalic (IT) = L6 Car3 neurons) compared to non-neurological controls for (columns from left to right) ALS-CI, ALS-nonCI, TDP-43 Pos (positive for TDP-43 pathology), and TDP-43 Neg (TDP-43 pathology negative) patients. The first panel represents all cryp-TE event genes from ALS-CI not in non-neurological controls, the second panel is the cryp-TE event genes shared between ALS-CI and TDP-43+5-Aza i3Neurons absent from non-neurological and i3Neuron controls, and the third and fourth panels show those shared genes which are significantly differentially up-(third panel) or down-regulated (fourth panel) in TDP-43+5-Aza vs control i3Neuron RNA-Seq. Each individual box (dark red with “*” = significant, FDR < 0.05) is colored by normalized enrichment score (NES) from Gene Set Enrichment Analysis (GSEA) of each gene set per panel for each differential gene expression set per group. Volcano plot (bottom panel) of fold change vs false discovery rate (FDR) q values for significantly differentially expressed genes only (dark grey) from snRNA-Seq BA46 L6 Car3 excitatory neurons in ALS-CI vs control patients. ALS-CI crypTE genes (events absent in control) detected by CrypTE-Seq are highlighted in dark red.

Overall, CrypTE-Seq allowed the detection of more than 2000 genes with cryptic TE splice sites per patient (**Fig 5B**, supplemental table **S14**). Looking globally at the relative fraction of all detected crypTE events, we saw similar distributions of crypTE-TSS, crypTE-exon, and crypTE-ApA transcripts as was detected in our i3Neurons (**Fig 4E** and **Fig 5B**). Furthermore, we noted a sharp increase in crypTE events in ALS-CI samples as compared to all other control and ALS/FTD tissues (**Fig 5B**), with the fewest crypTE transcripts captured in the ALS-nonCI samples that do not show TDP-43 pathology or cognitive involvement. Similarly to the i3Neurons, these crypTE events appear to occur at a basal level in control aged tissues, and increase in tissues with measured TDP-43 dysfunction.

To verify the TDP-43 dependence of these crypTE transcripts, we next compared the overlap between crypTE transcripts detected in the TDP-43+5-Aza i3Neurons and those captured in ALS/FTD patient tissues (**Fig 5C**). We first removed all crypTE transcripts that were detected in the CTRL i3Neurons or CTRL post-mortem BA46 tissues and then asked about shared crypTE events between TDP-43 depleted i3Neurons and the ALS/FTD cortical tissues (supplemental table **S15**). For crypTE-Exon events, 157 genes in ALS-CI, 90 in ALS-nonCI and 54 overlapping both ALS-CI and non-CI were identified to be shared with the TDP-43+5-Aza i3Neurons. In addition, the crypTE-Exon category for Fig 5C includes events which could be classified as either Exon or ApA, where we found 55 genes in ALS-CI, 30 in ALS-nonCI and 14 shared across all comparisons. For crypTE-ApA events, we detected the same crypTE transcripts for 34 gene-TE pairs present in ALS-CI samples, 16 in ALS-nonCI and 11 in both ALS-CI and ALS-nonCI, shared with i3Neurons. CrypTE-TSS events, which were difficult to detect in the ALS/FTD tissues, showed only 5 total crypTE fusion events shared between the i3Neurons and those detected in ALS/FTD tissues.

Next, we examined expression differences in genes which contain crypTE events in ALS-CI patients vs non-neurological controls. The volcano plots in **Fig 5D-E** show the Log2 fold change in gene expression from RNA-Seq data of TDP-43+5-Aza vs control i3Neurons on the horizontal axis and Log10 adjusted P-values (FDR) on the vertical axis with all significantly differentially expressed genes colored in dark grey. We have highlighted the genes with crypTE events specific to ALS-CI patients with: crypTE-Exon in orange (**Fig 5D)** and crypTE-ApA in yellow (**Fig 5E**). In total we saw significant TDP-43 dependent gene expression changes in i3Neurons for 466 genes with crypTE-Exon and/or crypTE-ApA events in ALS-CI patients (**Fig 5F**).

To further establish TDP-43 dependence of the ALS/FTD detected crypTE events, we incorporated our previously published TDP-43 eCLIP dataset from Tam et al. (2019)^20^. The TDP-43 eCLIP list enables us to determine the fraction of crypTE sequences known to be direct RNA binding targets of TDP-43. We found a significant enrichment between TDP-43 eCLIP targets and the genes with crypTE events in ALS-CI patient tissues (odds ratio = 1.73, p-value < 5.6e-20, Fisher’s exact test). This supports the TDP-43 dependent nature of these crypTE events found in ALS-CI patients and TDP-43 depleted i3Neurons, respectively.

We next focused on examples from the ALS/FTD patient data that had previously been seen in the TDP-43 depleted i3Neurons. Shown in **Fig 5G**, we detected the complex cassette inclusion event (crypTE-exon) that includes splicing from the sorting nexin gene SNX18 into two intronic TEs, LINE2c and LTR52. This event is present in the ALS-CI libraries, but not in non-diseased tissues or ALS tissues without evidence of TDP-43 dysfunction. Surprisingly, the ALS-nonCI tissues show mis-splicing into one of these intronic TEs (LTR-52) but not both, while tissues from patients with evidence of cognitive involvement and TDP-43 dysfunction show multiple mis-splicing events, including the complex event identified in i3Neurons.

Among the 34 crypTE-ApA events found to be shared between TDP-43 depleted i3Neurons and the ALS-CI patient tissues, we detected the NDEL1 event in both sets of samples (**Fig 5H**). This dynein transport complex gene shows clear signs of mis-splicing from exon 8 into a downstream LINE-2a element located approximately 30kb from the last annotated exon of NDEL1, as previously seen in TDP-43 depleted i3Neurons. This ApA event skips the last two exons of NDEL1 and substitutes LINE-2a sequence that would be included in the open reading frame (ORF) of the resulting fusion peptide. These shared crypTE events were detected in both the ALS cortical tissues and our i3Neuron *in vitro* model of TDP-43 dysfunction despite the fact that the bulk tissues samples contained a complex mixture of multiple neuronal and glial cell types.

### Genes that carry CrypTE isoforms show altered expression in layer-specific single neurons captured from ALS/FTD frontal cortex tissues with TDP-43 pathology and cognitive impairment

We next took advantage of single-nucleus transcriptome profiles (snRNA-Seq) of the prefrontal cortex region (n=24) from a larger cohort of patients that include these ALS-CI and ALS-nonCI donors^75^ (for more information on the cohort and methods please see Petrescu et al. ^75^). All donors were stratified by presence of TDP-43 pathology, clinical measures of cognitive impairment related to the BA46 region (executive function and working memory), and ALS diagnosis or age-matched non-neurological controls. By combining both snRNA-Seq and spatial transcriptomics, Petrescu et al. (2025) were able to categorise expression changes from layer specific excitatory and inhibitory neurons and glia. They showed that transcriptional alterations in deep layer (L5-L6) cortical neurons were the most strongly linked to measured alterations in cognitive impairment. Furthermore, genes with significant differential expression in ALS-CI patients vs control showed a significant enrichment for the same TDP-43 eCLIP targets from our (Tam et al, 2019) eCLIP gene list that we previously demonstrated was enriched among crypTE targets (**Fig 5F**). Therefore, in conjunction with data from our i3Neurons which model TDP-43 dysfunction in glutamatergic cortical-like excitatory neurons, we focused our comparisons on differentially expressed genes in the snRNA-Seq captured excitatory neurons from layers 4-6 of the BA46 region, where: layer 4 intratelencephalic excitatory neurons = L4, layer 5 extratelencephalic excitatory neurons = L5 ET, L5 intratelencephalic excitatory neurons = L5 IT, layer 6 corticothalamic excitatory neurons = L6 CT, and layer 6 Car3 marked intratelencephalic excitatory neurons = L6 Car3).

Comparing our list of crypTE targets discovered in the ALSci patient tissues to the ALSci nuc-seq profiles of Petrescu et al^75^, we found significant overlap in multiple excitatory neuron cell types (**Fig 5I**). In particular, we found the greatest enrichment for ALSci crypTE targets when looking at genes differentially expressed in ALSci L5-L6 excitatory neurons. In **Fig 5I**, we plot the normalized enrichment scores (NES) from a GSEA-based analysis of all differentially expressed genes in L4-L6 neuronal subtypes from ALS-CI, ALS-nonCI, TDP-43 positive and TDP-43 negative groups vs controls. In these GSEA calculations, the largest gene set used included all crypTE associated genes that were present in ALSci samples and absent in non-neurological controls **(Fig 5I**, blue vertical bar). We further stratified these ALSci crypTE genes by whether the crypTE event was shown to be TDP-43 dependent in i3Neurons (**Fig 5I**, pink vertical bar), and, in addition, whether those shared crypTE genes were also significantly up– or down-regulated in the i3Neurons upon TDP-43 depletion. A list of all GSEA calculated statistics is given in supplemental table **S16**.

To illustrate the concordance between crypTE genes and single-nucleus differential expression profiles underlying these GSEA calcutions, we’ve selected the cell type with strongest concordance between transcriptional alterations and presence of crypTE events: L6 Car3 IT neurons. The volcano plot in **Fig 5I** (lower panel) displays all genes with significant differential expression in ALS-CI vs control L6 Car3 IT excitatory neuron nuc-seq profiles (grey dots), with ALSci vs. control Log2 fold change on the horizontal axis and –Log10(FDR) on the vertical axis. Highlighted in dark red dots on the **Fig 5I** volcano plot, we identify all ALS-CI crypTE genes, where a strong bias toward up-regulation can be seen. This is of particular note as Petrescu et al. ^75^ found that L5 IT and L6 Car3 excitatory neurons differential gene expression profiles showed the strongest links to measures of cognitive impairment in their cohort. This suggests that crypTE splice alterations are contributing to the differential expression of genes linked to cognitive impairment in ALS.

## Discussion

Long-read RNA sequencing is a developing technology that enables the identification of full-length mature mRNA isoforms. This technology is greatly expanding our understanding of differential splicing^76^, long non-coding RNA species^77^, and expression from transposable elements (TEs)^78^. As this technology develops, it holds great promise for identifying disease-associated altered splicing events and novel mRNAs present across cellular types^79^. However, current long-read library preparation protocols show imperfect recovery of long, highly structured RNAs that are difficult to capture and amplify^80^. Here, we present optimized protocols for long-read RNA sequencing that improve on the ability to capture and retain full-length RNA transcripts, even from difficult to amplify species. This enabled the discovery of hundreds of TE transcripts that drive their own transcription, with locus-specific resolution on the origins of those TE reads. In addition to full-length expressed TEs, we were surprised to find many more examples of cryptic spliced gene-TE fusion transcripts.

TE exonization is a phenomenon in which cryptic splice sites within TE sequences are used as splice donor and acceptor sites that result in the inclusion of TE sequence within mature gene transcripts^81^ (crypTEs). These events are of interest as they provide a reservoir of novel gene and protein sequences that can alter gene regulation and function. While a few genes have been shown to regulate the TE exonization process^82,83^, this is the first study that links TDP-43 to TE exonization. We identified thousands of crypTE events that are normally prevented by TDP-43 protein in i3Neurons. We further demonstrate the presence of many of these TDP-43 dependent crypTE events in tissues from ALS/FTD patients with evidence of TDP-43 dysfunction. Given that these crypTE events are predicted by AlphaFold to include novel TE sequence within the resulting open reading frames, this suggests that crypTE events could be used as markers of TDP-43 dysfunction and hence expand the set of candidate TDP-43 biomarkers^41,44,84^. There is a severe need for biomarkers that can distinguish ALS and FTD patients from other neurodegenerative disorders as currently the diagnosis can only be confirmed upon autopsy. Current studies indicate that loss of nuclear TDP-43 may occur early in the disease process^85,86^, thus making TDP-43 dependent cryptic exons and crypTEs ideal candidates.

Our deep long-read RNA sequencing profiles enabled the detection of thousands of TDP-43 dependent crypTE transcripts present in cortical excitatory neuron models (i3Neurons) and also detected in postmortem ALS frontal cortex tissues from a cohort stratified by quantitative measures of cognitive impairment relevant to the brain region profiled. This enabled us to determine a set of crypTEs that may serve as reporters of cognitive dysfunction in ALS. We further incorporated single-nucleus transcriptome profiles to determine that these crypTEs were predominantly occuring in deep layer excitatory neurons (L6 Car3 IT) and may contribute to differential expression of those genes in ALS tissues. Finally, we provide a crypTE catalog that greatly expands the atlas of known TDP-43 dependent cryptic exons.

## Limitations of the Study

Long-read RNA sequencing studies generally involve mRNA capture techniques to avoid contamination from abundant ribosomal RNAs. Steps such as polyA capture can introduce bias in transcript coverage rates toward the 3’ end of gene regions. These polyA capture techniques limit the ability to reliably detect alterations at the 5’ end of transcripts as well as biasing against transcripts that do not contain extensive polyadenylation, such as histone genes^87^ and endogenous retroviruses (ERVs)^49^. Thus, we cannot rule out the possibility that we are missing a substantial number of crypTE-TSS events as well as expressed ERV sequences.

## Resource Availability

### Lead Contact

Further information and requests should be directed to and will be fulfilled by the lead contact, Molly Gale Hammell

## Materials availability

This study did not generate any new reagents; all reagents are commercially available.

## Data and code availability

Short read RNA sequencing data of i3Neurons is available at the following GEO accession: GSE310722. Long-read CrypTE-seq RNA libraries from i3Neurons and ALS cortical tissues are available at the following GEO accession: GSE310723 and GSE315972. Single-nucleus transcriptome profiles from ALS cortical tissues were obtained from Petrescu et al^75^ and are available at the following GEO accession: GSE290359.

## Supporting information

Supplemental Table Listing

## Acknowledgements

We would like to thank the M. Ward Lab, L. Gan Lab, Jackson Labs, and the NIH/iNDI consortium for the kind gift of i3Neuron cell lines. We would like to thank the M. Kampmann lab for gRNA expression vectors and advice on their use in i3Neuron lines. We thank RNAconnect for the kind gift of MarathonRT reagents and for helpful discussions. We would like to thank C. McLoughlin and M. Maurano for technical assistance with sequencing at NYU. We acknowledge the NYU LH Microscopy Lab and the NYU Genome Technology Center, which are partially supported by P30CA016087. PacBio sequencing was provided by the NYU Genome Technology Center (ULI TR001445). MGH was supported by NIH/NINDS (R01NS118570) and the Chan Zuckerberg Initiative’s Neurodegeneration Challenge Network.

## Author contributions

IB, RS, OHT, KN and MGH contributed to study design. IB, OHT, KO, CAJ, CGR and MGH contributed to data analysis and figure generation. RS and KN contributed to methodology. Experimental work was carried out by IB, RS, and KN. HP, CS, and the NYGC ALS Consortium provided patient tissue resources. All authors contributed to manuscript writing and approved the final manuscript.

## Methods

### Cell culture

The human induced pluripotent stem cells (iPSCs) were cultured, maintained, and cryopreserved as previously described (WC Skarnes et al, 2019). Briefly, iPSCs were maintained on Synthemax II-SC Substrate (Corning, Cat. no. 3535) coated tissue culture dishes in StemFlex complete medium (Thermo Fisher Scientific, Cat. no. A3349401) supplemented with Primocin (InvivoGen, Cat. no. ant-pm-05). Cultures were maintained at 37°C and 5% CO2, and the medium was replaced daily. Once confluent, cells were sub-cultured using ReLeSR (STEMCELL Technologies, Cat. no. 100-0483). For cryopreservation, cells were frozen using StemMACS Cryo-Brew (Miltenyi Biotec, Cat. no. 130-109-558). To thaw, cells were resuspended in StemFlex medium supplemented with 1X RevitaCell. All the cell culture work involving iPSC culture was approved by the Embryonic Stem Cell Research Oversight (ESCRO) committee and the Institutional Review Board (IRB) at the NYU Grossman School of Medicine.

### sgRNA Cloning and Lentiviral Packaging

Two non-targeting control (NT) sgRNA sequences were obtained from Shalem et al. (2014)^88^, and two sgRNA sequences targeting TDP-43 were obtained from Horlbeck et al. (2016)^89^. The sgRNAs against NT and TDP-43 were individually cloned into the pMK1334 lentiviral vector (Addgene, Cat. no. 127965) following the protocol described by Horlbeck et al. (2016). To generate high-titer lentiviral particles, the sgRNA plasmids were packaged in HEK-293T/17 cells (ATCC, Cat. no. CRL-11268) using Lenti-X™ Packaging Single Shots (VSV-G) (Takara Bio, Cat. no. 631276) according to the manufacturer’s protocol. The supernatant containing the virus was collected in sterile 50 ml tubes and concentrated using the Lenti-X™ Concentrator (Takara Bio, Cat. no. 631231) as per manufacturer’s protocol. The concentrated lentiviral aliquots were stored at –80°C until use.

### Cortical Neuron Differentiation and TDP-43 CRISPRi

A previously engineered CRISPRi-i^3^N-KOLF2.J1 iPSC line, obtained from The Jackson Laboratory (a gift from the NIH/INDi Consortium), was used for cortical neuron differentiation with minor modifications^33,36^. This line expresses human NGN2 under a doxycycline-inducible promoter (2 µg/mL) and constitutively expresses a dCas9-BFP-KRAB CRISPRi construct. Prior to differentiation, iPSCs received lentiviral particles encoding either non-targeting (NT) or TDP-43-targeting sgRNAs. The Lentiblast™ transduction enhancer (1:1000; OZ Biosciences, Cat. no. LBPX1500) was added to improve transduction efficiency. Forty-eight hours post-transduction, transduced cells were selected with 2 µg/mL puromycin (InvivoGen, Cat. no. ant-pr-1) for five days. The resulting stable cell pools were expanded before initiating differentiation. To initiate differentiation, 7 × 10 cells from the stable pools were dissociated using accutase (Innovative Cell Technologies, Inc., Cat. no. AT104), and seeded into AggreWell™800 plates (STEMCELL Technologies, Cat. no. 34815) using differentiation medium consisting of Neurobasal™ Plus Medium (Thermo Fisher Scientific, cat. no. A3582901), GlutaMAX™ (Thermo Fisher Scientific, cat. no. 35050061), BDNF (10 ng/mL; Thermo Fisher Scientific, cat. no. 450-02), NT-3 (10 ng/mL; Thermo Fisher Scientific, Cat. no. 450-03), doxycycline (2 µg/mL; Thermo Fisher Scientific, Cat.no.J67043.AD), and Gem21 NeuroPlex™ (without Vitamin A; Gemini Bio, Cat. no. 400-161-010), supplemented with 1X RevitaCell™ (Thermo Fisher Scientific, Cat. no. A2644501), to form spheroids^90^. After 24 hours, cell media was switched to fresh medium lacking RevitaCell, and changed daily. On day 8, neuronal spheroids were transferred and plated onto culture plates pre-coated with Geltrex™ (1:100; Thermo Fisher Scientific, Cat. no. A1413302) and Cultrex™ Stem Cell Qualified Laminin I (5 µg/mL; Bio-Techne, Cat. no. 3400-010-03). After overnight attachment, fresh medium was added, supplemented with 5 µM 5-Azacytidine, and the cells were incubated for 72 hours. Neuronal cultures were maintained in differentiation medium ± 5-azacytidine and harvested for analysis on either day 12 or day 15, depending on the specific downstream application.

### Immunofluorescence stains

For immunofluorescence analysis, cells were cultured in µ-Slide 4-Well chamber slides (ibidi, Cat. no. 80426), fixed with Fixation Buffer (BioLegend, Cat. no. 420801) for 15 minutes at room temperature, washed with PBS (no calcium and magnesium), and blocked with Intercept® (TBS) Blocking Buffer (LI-COR, Cat. no. 927-65001) for 2 hours at room temperature. The primary antibodies (TUBB3 (1:300; BioLegend, Cat. no. 801213), MAP2 (1:100; Thermo Fisher Scientific, Cat. no. PA5-17646), MAPT (Thermo Fisher Scientific, Cat. no. AHB0042) and TDP-43 (1:500; Proteintech, Cat. no. 10782-2-AP) were incubated overnight at 4°C. After washing, samples were incubated with Alexa Fluor-conjugated secondary antibodies (1:500; Thermo Fisher Scientific) for 2 hours at room temperature, and nuclei were counterstained with DAPI (1:500; Thermo Fisher Scientific, Cat. no. 62248). Stained slides were mounted using ProLong™ Diamond Antifade Mountant (Thermo Fisher Scientific, Cat. no. P36965). Mounted slides were scanned using a Zeiss LSM 880 confocal microscope with 40X and 63X objectives, and all subsequent image analysis and figure generation were performed using ImageJ software^91^.

### RNA-seq library generation

RNA was extracted and purified from cultured cells using the Direct-zol™ RNA Miniprep Plus (R2071, Zymo Research, Irvine, CA, USA). RNA concentration was determined with the Qubit RNA BR Assay Kit (Q10210, ThermoFisher Scientific, Waltham, MA, USA) and Qubit 4 Fluorometer (Q33226, ThermoFisher Scientific, Waltham, MA, USA). RNA quality was assessed using the RNA 6000 Nano (5067-1511, Agilent, Santa Clara, CA, USA) on the Agilent 2100 Bioanalyzer (G2939BA, Agilent, Santa Clara, CA, USA). RNA-seq libraries were prepared from 1000 ng of total RNA subjected to rRNA depletion with the NEBNext rRNA Depletion Kit v2 (Human/Mouse/Rat) (E7405L, New England Biosciences, Ipswich, MA, USA). Strand specific library preparation was performed using the NEBNext II Ultra Directional Library Prep Kit (E7760S, New England Biosciences, Ipswich, MA, USA) and sequenced on the NextSeq 500 using a single end 75 nucleotide setting, to yield approximately 50-60 million reads per library.

### RNA-seq differential expression and alternate splicing analysis

Reads were aligned to the hg38 human genome using STAR v2.7.6^92^, allowing up to 100 alignments per read to capture transposon sequences. Abundance of gene and transposon reads was calculated with TEtranscripts v2.2.3^93^. For differential expression analysis, we used DESeq2^94^ with DESeq’s normalization strategy and negative binomial modeling. Comparisons with a Benjamini-Hochberg corrected P-value of p < 0.05 were considered significant. Gene expression estimates were normalized using a variance stabilizing transformation in DESeq2 for heatmap visualization. Alternative splicing was performed using Leafcutter^43^ following the procedure as outlined in their documentation (http://davidaknowles.github.io/leafcutter/articles/Usage.html)

### RNA isolation, cDNA synthesis and RT-qPCR analysis

Total RNA was isolated from cortical neurons in triplicate for each experimental condition using the Direct-zol RNA Miniprep Plus Kit (Zymo Research, Cat. no. R2071) according to the manufacturer’s instructions. For each sample, 500 ng of the isolated RNA was used to synthesize cDNA with the SuperScript VILO cDNA Synthesis Kit (Thermo Fisher Scientific, Cat. no. 11754050), following the manufacturer’s protocol.

Pre-designed PrimeTime qPCR primers for TDP-43 (IDT, Cat. no. Hs.PT.58.26912658) and the housekeeping gene RPL13A (IDT, Cat. no. Hs.PT.58.45725862) were obtained from Integrated DNA Technologies. Quantitative real-time PCR (RT-qPCR) was performed in triplicate using PowerUp SYBR Green Master Mix (Thermo Fisher Scientific, Cat. no. A25742). The polymerase reactions were run on a QuantStudio 7 Real-Time PCR System (Thermo Fisher Scientific). The relative expression values were calculated using the ΔΔCt method^95^.

### CrypTE-seq long read RNA sequencing

RNA quality was evaluated using the RNA 6000 Nano (5067-1511, Agilent, Santa Clara, CA, USA) kit run on the Agilent 2100 Bioanalyzer (G2939BA, Agilent, Santa Clara, CA, USA). CrypTE-seq read libraries were prepared from ∼300ng total RNA per sample following a modified PacBio Iso-Seq v2 library protocol (103-071-500, PacBio, Menlo Park, CA, USA) and pooled using the SMRTbell Prep Kit 3.0 (102-182-700, PacBio, Menlo Park, CA, USA). Modifications include the use of post-primer annealing with first strand cDNA synthesis performed with MarathonRT (200035, RNAConnect, Branford, CT, USA) followed by Iso-seq RT and then addition of the template switch oligo. All other steps followed the Iso-seq v2 library protocol (103-071-500, PacBio, Menlo Park, CA, USA). Samples were loaded on a Revio SMRT cell tray using the Revio Polymerase loading kit (102-739-100, PacBio) at 500pM concentration and sequenced on a PacBio Revio long read sequencer using the PacBio application Iso-Seq Method with Standard library.

### Bioinformatic processing of long read RNA sequencing

Circular consensus sequencing reads were generated from PacBio sequencing output using the CCS algorithm (version 8, Pacific Biosciences). Libraries were demultiplexed, refined and clustered using IsoSeq toolkit (version 4.0.0). FASTA files were generated and aligned to the hg38 genome using minimap2^96^ (version 2.24) with splice:hq preset (−x splice:hq) and GT-AG search on the transcript strand (−u f). Transcript discovery and quantification was performed using bambu^97^ (version 3.2.6), with the GENCODE^98^ gene annotation (version 47) as reference.

### Identification and characterization of gene-TE fusion transcripts

Aligned transcripts were annotated against GENCODE gene annotations (version 47) and UCSC RepeatMasker^99^ annotations for the hg38 genome. Transposable elements were classified as having an intact 5’ end if the annotation starts at the first nucleotide (for Alu, LTR and SVA) or before nucleotide 150 (for young L1) of the consensus sequence. Transcripts containing non-overlapping gene exonic and transposable element annotations were extracted. Chimeric transcripts are classified as CrypTE-TSS if the TE is within 50 bp of the transcript start position, CrypTE-ApA if the TE is within 50 bp of the transcript end position, or CrypTE-exon if it is >50 bp from either end of the transcript. CrypTE-exon events were further filtered to ensure that the splice donor/acceptor sites occur within the TE sequence to exclude simple read-through transcription. Open reading frames (ORF) are determined using a custom perl script, and ORFs >=1kb (or longest ORF if it is <1kb) are assessed for potential nonsense mediated decay using a custom perl script based on the 50-nt rule^58^. Chimeric protein structures were predicted using AlphaFold^59^ 3 model (alphafoldserver.com). Spliced junctions of gene-annotation-only transcripts were compared against GENCODE (version 47), and were classified to be: “novel splicing” if the cryptic splice junction joined two known exons of the same gene; “novel-exon” if the cryptic splice junction joined to an unannotated genomic region; and “gene fusion” if the cryptic splice junction joined two exons of different genes.

### Tissue sourcing and donor information

Postmortem brain tissue was sourced from the Medical Research Council (MRC) Edinburgh Brain Bank (Research Ethics Committee approval: 21/ES/0087) with appropriate ethical approvals. Samples were obtained from ALS patients enrolled in the Scottish Motor Neurone Disease Register who had consented to collection and research use of their health-related data. The clinical metadata (including ECAS test scores) was sourced from the Clinical Audit Research Evaluation (CARE)-MDN database for each patient. The Edinburgh Sudden Death Brain Bank^100^ provided tissue from neurotypical control cases (donors without neurological or psychiatric disorders and lacked significant post-mortem neuropathology). All material transfers complied with applicable federal, state and local regulations. ALS diagnoses met the revised El Escorial criteria for clinically definite or probable disease. Individuals with comorbid neurological, including frontotemporal dementia, or psychiatric conditions were excluded, as were donors with clinically documented dementia.

### Defining sharing of crypTE events between ALS-CI and/or ALS-nonCI with i3Neurons

To focus on shared crypTE events between either or both ALS-CI and –nonCI patients with TDP-43+5-Aza i3Neurons, if the exact crypTE-gene event was detected in comparisons with control for ALS-CI vs non-neurological controls, ALS-nonCI vs non-neurological controls, and TDP-43+5-Aza vs control i3Neurons, then the crypTE transcripts was filtered to exclude these entries. Then those events remaining were compared across condition and events with the same crypTE-gene pairing with identical TE copies involved were defined as shared between either/both ALS-CI/ALS-nonCI and TDP-43+5-Aza i3Neurons. Events were then filtered by crypTE event category: crypTE-TSS, –Exon, –ApA, –TSS/Exon, –TSS/ApA, –Exon/ApA and –TSS/Exon/ApA. The count for a cryTE containing gene with a shared event was assigned separately per crypTE event type and if that gene had multiple shared crypTE-gene events with TDP-43+5-Aza i3Neurons, those duplicates were collapsed into one count for that gene if all were from only either ALS-CI or ALS-nonCI. To address events which could be classified as more than one type of crypTE event, if that gene also had a shared crypTE event of another single category (−TSS, –Exon or –ApA) for the same group (ALS-CI, ALS-nonCI or both) then that gene was reassigned to that category and handled as above. In cases where there were additionally counts for sharing between i3N TDP-43+5-Aza i3Neurons with both ALS-CI and ALS-nonCI, then that gene was assigned only to the category of “shared with i3N TDP-43+5-Aza: ALS-CI + ALS-nonCI” and not counted as part of “ALS-CI” or “ALS-nonCI” for that crypTE event type.

---

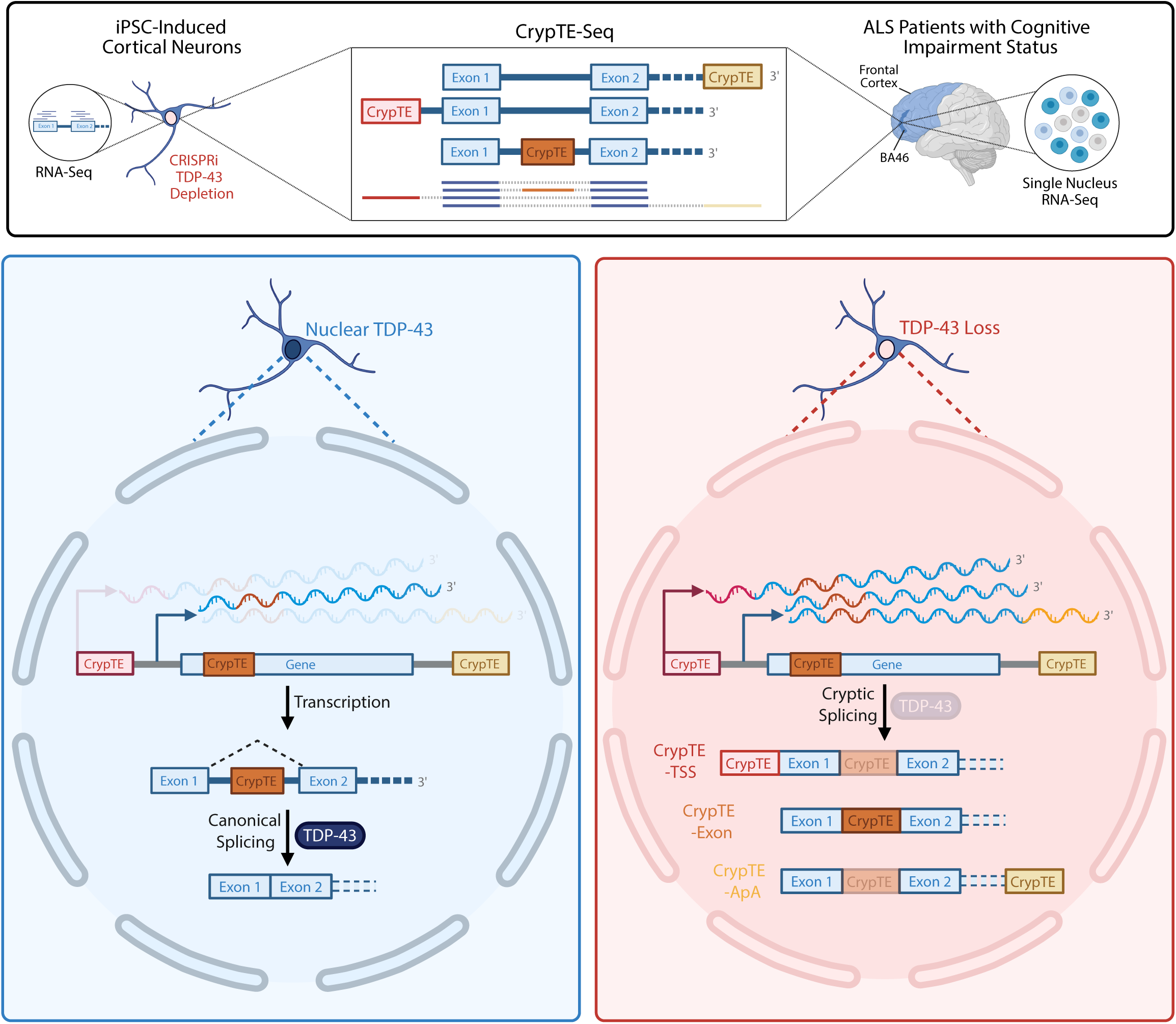

