## Supplemental Table Listing for "TDP-43 dysfunction leads to the accumulation of cryptic transposable element-derived exons, crypTEs, in iPSC derived neurons and ALS/FTD patient tissues"

**Supplementary Table Legends**

**Supplemental table S1A.** Differential gene expression in TDP-43+5-Aza vs control i3Neurons.

**Supplemental table S1B.** Differential gene expression in TDP-43 KD vs control i3Neurons.

**Supplemental table S1C.** Differential gene expression in 5-Aza vs control i3Neurons.

**Supplemental table S2A.** Annotated leafcutter output of TDP-43+5-Aza vs control i3Neurons.

**Supplemental table S2B.** Annotated leafcutter output of TDP-43 KD vs control i3Neurons.

**Supplemental table S2C**. Annotated leafcutter output of 5-Aza vs control i3Neurons.

**Supplemental table S3.** Delta percent spliced in (PSI) for differential spliced junctions in Liu et al. 2019 and i3Neurons from this study.

**Supplemental table S4A.** Summary of detected transcripts in i3Neuron CrypTE-Seq libraries.

**Supplemental table S4B.** Annotated Bambu output of i3Neuron CrypTE-Seq libraries.

**Supplemental table S5.** Overlap between differentially spliced junctions identified in Ma et al. (2022) and Seddighi et al. (2024) with CrypTE-Seq identified differential isoform usage in i3Neurons from this study.

**Supplemental table S6A.** TE transcripts identified with CrypTE-Seq in TDP-43+5-Aza i3Neurons.

**Supplemental table S6B.** TE transcripts identified with CrypTE-Seq in TDP-43 KD i3Neurons.

**Supplemental table S6C.** TE transcripts identified with CrypTE-Seq in 5-Aza i3Neurons.

**Supplemental table S6D.** TE transcripts identified with CrypTE-Seq in control i3Neurons.

**Supplemental table S7A.** CrypTE transcripts identified in TDP-43+5-Aza i3Neurons with CrypTE-Seq.

**Supplemental table S7B.** CrypTE transcripts identified in TDP-43 KD i3Neurons with CrypTE-Seq.

**Supplemental table S7C.** CrypTE transcripts identified in 5-Aza i3Neurons with CrypTE-Seq.

**Supplemental table S7D.** CrypTE transcripts identified in control i3Neurons with CrypTE-Seq.

**Supplemental table S8A.** Cryptic non-TE transcripts identified in TDP-43+5-Aza i3Neurons with CrypTE-Seq.

**Supplemental table S8B.** Cryptic non-TE transcripts identified in TDP-43 KD i3Neurons with CrypTE-Seq.

**Supplemental table S8C.** Cryptic non-TE transcripts identified in 5-Aza i3Neurons with CrypTE-Seq.

**Supplemental table S8D.** Cryptic non-TE transcripts identified in control i3Neurons with CrypTE-Seq.

**Supplemental table S9A.** non-TE gene fusion transcripts identified in TDP-43+5-Aza i3Neurons with CrypTE-Seq.

**Supplemental table S9B.** non-TE gene fusion transcripts identified in TDP-43 KD i3Neurons with CrypTE-Seq.

**Supplemental table S9C.** non-TE gene fusion transcripts identified in 5-Aza i3Neurons with CrypTE-Seq.

**Supplemental table S9D.** non-TE gene fusion transcripts identified in control i3Neurons with CrypTE-Seq.

**Supplemental table S10.** NMD Prediction for crypTE transcripts in TDP-43+5-Aza i3Neurons not in control.

**Supplemental table S11.** Predicted crypTE fusion proteins in TDP-43+5-Aza i3Neurons not in control.

**Supplemental table S12.** Metadata of ALS/FTD patient samples used for crypTE analysis

**Supplemental table S13A.** Summary of detected transcripts in Patient CrypTE-Seq libraries.

**Supplemental table S13B.** Annotated Bambu output of Patient CrypTE-Seq libraries

**Supplemental table S13C**. Table S13C: Overlap between differentially spliced junctions identified in Ma et al. (2022) and Seddighi et al. (2024) with patient isoseq study.

**Supplemental table S14A.** CrypTE transcripts detected in ALS-C9-03363 (CI-1) BA46 sample.

**Supplemental table S14B.** CrypTE transcripts detected in ALS-CI-02988 (CI-2) BA46 sample.

**Supplemental table S14C.** CrypTE transcripts detected in ALS-CI-04118 (CI-3) BA46 sample.

**Supplemental table S14D.** CrypTE transcripts detected in ALS-C9-02759 (nonCI-1) BA46 sample.

**Supplemental table S14E.** CrypTE transcripts detected in ALS-02763 (nonCI-2) BA46 sample.

**Supplemental table S14F.** CrypTE transcripts detected in ALS-03634 (nonCI-3) BA46 sample.

**Supplemental table S14G.** CrypTE transcripts detected in CTL-03662 (CTRL) BA46 sample.

**Supplemental table S14H.** CrypTE transcripts detected in CTL-03374 (CTRL) BA46 sample.

**Supplemental table S15.** Shared crypTE-TSS, -Exon and -ApA events between ALS/FTD Patients and TDP-43+5-Aza i3Neurons not in Control i3Neurons or Non-neurological Control BA46 Post-mortem Tissue.

**Supplemental table S16.** GSEA results for ALS-CI crypTE genes (all, shared with TDP-43+5-Aza i3Neuron CrypTE Genes and/or up-/down-regulated in TDP-43+5-Aza i3Neurons) enriched in snRNA-Seq differentially expressed genes for ALS-CI, ALS-nonCI, TDP-43 pos and TDP-43 neg patients compared to control.
